# Associations between Early Midlife Lifestyle Behaviors, Young Adult Cognitive Reserve and Advanced Predicted Brain Age in Late Midlife

**DOI:** 10.1101/2020.11.02.362780

**Authors:** Carol E. Franz, Sean N. Hatton, Jeremy A. Elman, Teresa Warren, Nathan A. Gillespie, Nathan A. Whitsel, Olivia K. Puckett, Anders M. Dale, Lisa T. Eyler, Christine Fennema-Notestine, Donald J. Hagler, Richard L. Hauger, Ruth McKenzie, Michael C. Neale, Matthew S. Panizzon, Rahul C. Pearce, Chandra A. Reynolds, Mark Sanderson-Cimino, Rosemary Toomey, Xin M. Tu, Hong Xian, Michael J. Lyons, William S. Kremen

## Abstract

**Importance:** Both cognitive reserve and modifiable lifestyle behaviors are associated with dementia risk. The effect of early lifestyle behaviors and cognitive reserve on late midlife brain aging could inform early identification and risk reduction of future dementia.

**Objective:** Determine associations of young adult cognitive reserve, early midlife lifestyle behaviors, and the reserve-by-lifestyle interaction on late midlife brain age. Examine the relationship between mild cognitive impairment (MCI) and brain age.

**Design:** Participants were from the nationally representative Vietnam Era Twin Study of Aging (VETSA). Cognitive reserve was assessed at median age 20 years (IQR=1.38) with the Armed Forces Qualification Test (AFQT). Lifestyle behaviors (smoking, alcohol consumption, and social engagement) were assessed at median age 41 (IQR=5.00). Structural brain imaging conducted at median age 69 (IQR=4.74) was used to construct predicted brain age difference scores (PBAD=chronological age minus predicted brain age) and MCI was ascertained.

**Setting:** In-person cognitive testing (ages 20 and 69); mailed survey (age 41); structural MRI, MCI diagnosis at University of California, San Diego (age 69).

**Participants:** 431 community-dwelling men.

**Exposures:** AFQT; self-reported health and lifestyle behaviors.

**Main outcomes and measures:** PBAD scores; MCI.

**Results:** In fully adjusted mixed linear models, age 20 cognitive reserve and the age 41 lifestyle composite were associated with age 69 PBAD [t (104)=2.62, p=0.01, 95%CI 0.874, 6.285; t (104)=3.37, p=0.001, 95%CI 0.583, 2.249 respectively] as was the reserve-by-lifestyle interaction [t (104) = −2.29, p=0.02, 95%CI −2.330, −0.167]. Unfavorable lifestyle predicted more advanced brain age, but only for those with lower young adult cognitive reserve. The MCI group had more advanced brain age (F (2,130) = 3.13; p=0.05).

**Conclusions and relevance:** Favorable lifestyle behaviors promoted resistance to accelerated brain aging 3 decades later for those with lower young adult cognitive reserve. High reserve appeared to be protective regardless of lifestyle. The association with MCI suggests that advanced PBAD scores reflect poorer brain integrity, although it is unclear if PBAD is related to Alzheimer’s dementia specifically. Lower cognitive reserve increases risk for dementia, but early lifestyle modification may promote healthier brain aging and dementia risk reduction, particularly in those with lower reserve.

**Study Type:** Cohort Study

**Key Points:** *Question:* Do modifiable lifestyle behaviors in early midlife predict later accelerated brain aging and is that association moderated by cognitive reserve?

*Findings:* A lifestyle composite of smoking, alcohol consumption and social engagement at age 41 was associated with estimated brain age in late midlife. There was a significant moderation effect whereby more unfavorable lifestyle behaviors predicted more advanced brain aging, but only in those with low-to-moderate cognitive reserve.

*Meaning:* Favorable lifestyle behaviors appear to be protective for brain integrity especially among those with lower cognitive reserve. Early midlife efforts at prevention could be prioritized among those with lower cognitive reserve.

## Introduction

With increasing numbers of individuals developing mild cognitive impairment (MCI) and Alzheimer’s Disease (AD), early risk reduction and prevention efforts are viewed as important in reducing and delaying dementia onset.^1–3^ Major guidelines and reviews consistently associate modifiable lifestyle behaviors of physical activity, tobacco use, diet, alcohol consumption, and social engagement with dementia risk.^1–9^ The 2020 Lancet Commission report proposed that modification of lifestyle behaviors could reduce dementia incidence by as much as 35%.^1–2^ Higher cognitive reserve—an individual’s overall cognitive capacity—is also protective against decline and dementia.^10–13^ In recent years, the concept of biomarkers of “biological age” has gained traction^14,15^ and has been applied to models of brain aging.^16,17^ Advanced brain aging has been observed in both MCI and AD, and may serve as a risk indictor for dementia.^18,19^ Toward the goal of early intervention and risk reduction, the primary aim of this project was to investigate associations of early midlife lifestyle behaviors and young adult cognitive reserve, including a lifestyle-by-reserve interaction, with late midlife brain aging.

Machine learning algorithms that predict chronological age in large samples of typically aging adults are applied to magnetic resonance imaging (MRI) features to calculate predicted brain age, a global measure of brain integrity.^17,18,20^ Advanced or accelerated brain aging is inferred when predicted brain age is older than expected given one’s chronological age. Cognitive decline from childhood has been associated with advanced brain age in midlife.^20,21^ AD patients’ brain age appeared older than their chronological age by about 10 years^18^ and brain age was more advanced in MCI cases who progressed to AD than those who did not.^19^ Among AD and MCI cases who progressed to AD, 3-year longitudinal increases in predicted brain age were greater in *APOE-ε4* carriers than non-carriers.^22^

Lifestyle findings consistently support a dose-dependent response in that co-occurrence of multiple risky lifestyle behaviors is associated with worse outcomes compared with single risks.^5,9^ Among adults ages 55-85, for example, a favorable lifestyle composite was associated with healthier brain characteristics.^23^ To date, several studies using predicted brain age measures have examined individual lifestyle behaviors, but few have examined lifestyle composites.^17,24^ Concurrent associations of older brain age with smoking and alcohol consumption, but not physical activity, have been observed in middle-aged adults.^17,24,25^ The effect of alcohol dependence on predicted brain age was much stronger in older adults (as much as 11.7 years older) compared with middle aged adults.^26^ Brain age was also significantly correlated with type 2 diabetes, smoking duration, alcohol consumption, and depression within-time in adults in their mid-60s.^16,27^ Adults who both drank and smoked had more advanced brain age than those with only one of these behaviors.^28^ These cross-sectional results suggest that favorable lifestyle factors may promote resistance to accelerated brain aging. Theoretically, having a younger, healthier brain at midlife could ultimately reduce or delay onset of MCI or AD and other dementias.^13^

Despite the documented importance of cognitive reserve^13^ studies with direct measures of early cognitive reserve are rare. However, these are important because they help to address the issue of reverse causation. Age 11 childhood cognitive reserve, controlling for smoking and socioeconomic status, was a risk factor for vascular dementia but not AD in a 1921 Scottish birth cohort^10^ and higher age 11 cognitive reserve was associated with thicker cortex at age 73 in a 1936 Scottish birth cohort.^11^ Worse cognitive and brain health at age three predicted more advanced brain age in adults at age 45, and older brain age was correlated with signs of advanced aging such as poorer health and older facial features.^21^ Cognitive reserve, then, may also promote resistance to accelerated brain aging. However, to our knowledge, there are no studies examining lifestyle-by-cognitive reserve interactions on brain age.

In the present study, we hypothesized that having fewer favorable lifestyle behaviors or lower young adult cognitive reserve would be associated with more advanced brain aging. We also tested a lifestyle-by-cognitive reserve interaction and predicted that young adult cognitive reserve would moderate the association between lifestyle behaviors and brain age. Finally, we examined whether predicted brain age is an indicator of cognitive decline by evaluating associations between MCI and brain aging in late midlife.

## Methods

### Participants

Participants were community-dwelling non-patient men from the Vietnam Era Twin Study of Aging (VETSA).^29^ VETSA comprise a sample of male-male twins who are members of the nationally representative Vietnam Era Twin (VET) Registry. They are similar to American men in their age range with respect to health and lifestyle characteristics.^30^ The majority (~80%) reported no combat exposure. VETSA participants were randomly recruited from 3322 twin pairs in the VET Registry.^31^ Ages at the three different assessments are provided below. Average lifetime education was 13.98 (SD 2.07) years.

### Procedures

#### Young Adult Cognitive Reserve (median age 20)

Participants completed the Armed Forces Qualification Test (AFQT) in person (median age=19.51 years; IQR=1.38; range=17-25; 1965-1975) during induction into the military. The AFQT is a standardized, validated 100-item multiple-choice paper-and-pencil test of general cognitive ability (median age=19.51 years; IQR=1.38; range=17-25).^32^ This test is highly correlated with other tests of GCA such as Wechsler Adult Intelligence Scale (*r*=0.84).^33^ Average intelligence of participants was estimated at 105.^33^

#### Early Midlife Modifiable Lifestyle Behaviors (median age 41)

In 1990, in early midlife (median age=41; IQR=5.00; range=34-44), participants completed a mailed health survey which queried their smoking, alcohol consumption, social engagement, physical activity and diet behaviors (see Supplemental eTable 1).^34^ Each modifiable lifestyle indicator was operationalized based on standard criteria and coded as favorable (1) or unfavorable (0).^7,8^

Smoking status was defined as current (unfavorable) versus never/former (favorable). Unfavorable alcohol consumption was defined as drinking >2 drinks per day across the past 14 days. Drinking ≤2 drinks per day, former, or never drinking was scored as favorable. Social engagement was operationalized based on 6 measures: marriage (currently married/not married), involvement in church more than twice a month (yes/no), clubs/organizations (yes/no), number of close friends (≥3;<3), number of close relatives (≥3;<3), and number of close people connected with at least once a month (≥3;<3).^7^ Social engagement scores of 0-2 were categorized as unfavorable and 3-6 as favorable.

Favorable physical activity reflected engaging in moderate-to-vigorous activity at least 3 days per week; less activity was coded as unfavorable. Favorable diet was coded if a participant met recommendations for 3 of 5 food groups per week: fish, fruits and vegetables, dairy, red meat, and processed meat. Table 1 displays the proportion of favorable classifications for each indicator. Frequencies of the 5 modifiable lifestyle behaviors at age 41 ranged from a low of 53% who met criteria for a favorable diet to 87% who met criteria for favorable alcohol consumption.

**Table 1.** Early Midlife Favorable Lifestyle Behaviors and Frequencies.

| Behavior | Definition of Favorable rating<br>(0=unfavorable/1=favorable) | N (%) Favorable<br>Lifestyle |
| --- | --- | --- |
| Alcohol Consumption | Drinks $\leq 2$ drinks containing alcohol per day | 409 (87.4%) |
| Tobacco Smoking | Never or former | 342 (72.6%) |
| Social Engagement | Engages in 3-6 forms of social interaction regularly | 360 (77.8%) |
| Diet | Consumes 3-5 Healthy food groups per day | 243 (53.3%) |
| Physical activity | Moderate to vigorous activity at least 3 days/week | 386 (83.4%) |
| Lifestyle Composite | Sum of alcohol consumption, smoking and social engagement; higher score=more favorable behaviors | 0 = 6 (1.3%)<br>1 = 50 (11.2%)<br>2 = 165 (36.9%)<br>3 = 226 (50.6%) |

#### Late Midlife Data Collection (median age 69)

In-person data collection during late midlife (median age=68.90 years; IQR=4.74; range= 61-72; 2016-2019), included a detailed medical history interview, socio-demographic data, in-depth cognitive testing, and structural MRI for those who passed a standard MRI safety screening (N=481). More details on VETSA procedures are published elsewhere.^35–37^

##### MRI Acquisition and Processing (median age 69)

T1-weighted images providing high anatomical detail were acquired on 2 GE 3T Discovery 750× scanners (GE Healthcare, Waukesha, WI, USA) with an 8-channel phased array head coil (scanner 1 N=336, scanner 2 N=145) at the University of California, San Diego. The T1 imaging protocol included a sagittal 3D fast-spoiled gradient echo T1-weighted image (echo time=3.164 msec, repetition time=8.084 msec, inversion time=600 msec, flip angle=8°, pixel bandwidth=244.141, field of view=25.6 cm, frequency=256, phase=192, slices=172, slice thickness=1.2 mm). As described in prior work raw image files were processed using an in-house pipeline by the UCSD Center for Multimodal Imaging and Genetics.^38^ Data were qualitatively assessed and images with severe scanner artifacts or excessive head motion were rescanned where possible or excluded from analysis (~3%). T1-weighted structural images were corrected for gradient distortions^39^ and B1 field inhomogeneity.^40^ Subcortical segmentation and surface-based cortical parcellation were performed using FreeSurfer version 5.1 (surfer.nmr.mgh.harvard.edu) as previously described.^41^ Inaccuracies in automated segmentations were manually corrected by trained neuroimaging analysts. All images required some form of manual editing to ensure the correct classification of the pial and white matter surfaces, with particular attention given to the orbitofrontal cortex, the temporal lobes, meninges, and transverse and superior sagittal sinuses. Problematic segmentations/parcellations were reviewed for exclusion by consensus with four neuroimaging analysts.

##### Late Midlife Predicted Brain Age

We used the Brain-Age Regression Analysis and Computation Utility software BARACUS v0.9.4^20^ (https://github.com/BIDS-Apps/BARACUS; https://zenodo.org/record/826543#.WjF5Ft-nE2x). BARACUS uses linear support vector regression models to predict brain age in adults derived from each individual’s FreeSurfer statistics.^15^ Specifically, vertex-wise cortical metrics were derived from the fsaverage4 standard space for cortical thickness (n=5124 vertices) and surface area (n=5124 vertices), and subcortical segmentation metrics were derived from the aseg.stat file for subcortical volume (n=66 regions of interest). We used the BIDS-mode docker on Ubuntu 16.04 using the default database (Liem2016_OCI_norm), which is trained on 1166 independent subjects with no objective cognitive impairment (566 female/600 male, 20-80 years).^20^ The predicted brain age difference score (PBAD) is calculated by subtracting predicted brain age (referred to as “stacked-anatomy” brain age in BARACUS) from chronological age. Therefore, a negative PBAD is indicative of brain age that is estimated to be older than one’s chronological age. PBAD was adjusted, via regression, for scanner. One participant was excluded due to brain cancer and 2 due to brain damage of unknown origin.

##### Late Midlife Mild Cognitive Impairment (MCI)

The Jak-Bondi approach was used to diagnose MCI^42–44^ based on 18 cognitive tests covering memory, executive function, attention, language, visuospatial ability, and processing speed.^29^ Criteria for impairment within a domain required performance on 2+ tests that were each >1.5 SDs below age- and education-adjusted normative means. Cognitive scores were adjusted for young adult GCA in order to differentiate decline from lifelong poor performance.^45^ Scores were additionally adjusted for practice and attrition effects using the replacement-subjects method as described previously.^46^

### Covariates

Ethnicity was defined as non-Hispanic White versus other. Early midlife (age 41) covariates included a cardiovascular index: total of yes/no responses to angina pectoris, congestive heart failure, coronary heart disease, damaged heart valves, heart attack/myocardial infarction, phlebitis/thrombophlebitis, and stroke. The respiratory index included asthma, chronic obstructive pulmonary disease, and emphysema. Liver disease index comprised alcoholic hepatitis, cirrhosis of the liver. One item addressed physician diagnosed depression (yes/no). Due to the low frequency of health problems, the final cardiovascular, respiratory, and liver disease indices are categorized as any disorders versus none. Hypertension and diabetes were coded based on the response to “Have you ever been told by a doctor that you had —.” Obesity was coded as BMI ≥30 based on self-reported height and weight. Self-reported stroke at age 69 was also included as a covariate. Frequencies of these health items and their correlations with key variables are shown in Supplemental eTables 2 and 3. *APOE* genotype was classified as having ε4+ versus ε4-.^47^ Some data were missing for *APOE* (N=63) since not all participants provided blood or were genotyped. Participants were excluded if they self-reported a history of seizures, multiple sclerosis, HIV, or schizophrenia (N=6), leaving a final N of 431 with complete PBAD, cognitive reserve, lifestyle, and demographic data.

### Statistical analysis

In preliminary analyses, alcohol consumption, smoking, and social engagement were the only lifestyle behaviors associated with later PBAD at p<.10 (Supplemental eTable 4). These 3 measures were summed to create a lifestyle composite (0-3). As seen in Table 1, 51% of the sample met criteria for favorable ratings on all 3 behaviors. Table 2 shows associations between key variables and number of favorable lifestyle behaviors.

**Table 2.** Basic Demographics and Descriptive Statistics.

|  | Full Sample | Number Favorable Lifestyle Behaviors (Composite) |  |  |  | F test/p value |
| --- | --- | --- | --- | --- | --- | --- |
|  |  | 0 (N=5) | 1 (N=50) | 2 (N=164) | 3 (N=224) |  |
| Age @ Lifestyle assessment: mean (SD) | 39.95 (2.68); Range 33-44 | 38.40 (2.19) | 38.88 (2.60) | 39.69 (2.74) | 40.45 (2.57) | F (3,147)=1.22<br>p=0.35 |
| Age @ MRI scan: mean (sd) | 67.61 (2.62); Range 61-72 | 65.94 (2.80) | 66.48 (2.56) | 67.44 (2.67) | 68.05 (2.46) | F (3,147)=0.37,<br>p=0.78 |
| Young Adult Cognitive Reserve: mean (SE) | 0.33 (0.67) | 0.52 (0.46) | 0.14 (0.70) | 0.25 (0.70) | 0.48 (0.63) | F (3,144)=2.55,<br>p=0.058; 2<3 |
| Brain age difference score: mean (SE) | 0.82 (0.38) | -3.63 (2.19) | -2.31 (0.74) | -.009 (0.42) | 0.82 (0.38) | F (3,147)=6.11<br>p=.0006; 0<3; 1<2,3 |
| Ethnicity: N (%) non-Hispanic White | 398 (90%) | 5 (100%) | 42 (84%) | 143 (87.2%) | 208 (92.7%) | $\chi^2(3,443)=0.12$, ns |
| Stroke: N (%) at VETSA 3 | 28 (5.4%) | 1 (20%) | 6 (12%) | 7 (4.3%) | 4 (1.8%) | $\chi^2(3,443)=14.34$,<br>p=.003 |
| *APOE*: N (%) no e4 allele | 285 (75%) | 3 (60%) | 33 (76.7%) | 100 (74%) | 149 (75.6%) | $\chi^2(3,380)=0.86$, ns |

Table note: For F tests, N=range from 431 (age 20 AFQT) to 437 (others) but df is lower due to random effect of case.

We conducted linear mixed models in SAS 9.4 using maximum likelihood methods. The basic multivariable model (Model 1) examined the main effects of young adult cognitive reserve and the early midlife lifestyle composite, as well as the cognitive reserve-by-lifestyle interaction on late midlife PBAD adjusting for age, ethnicity, and stroke. Analyses treated twins as individuals so familial relationship was accounted for by including family as a random effect. The full model (Model 2) additionally adjusted for early midlife cardiovascular problems, hypertension, obesity, diabetes, respiratory problems, liver disease, depression, *APOE* genotype, and height. The N in the full model is 354, primarily because of missing *APOE* data. All results are reported as Type III tests of fixed effects, two-tailed.

## Results

In the basic multivariable model (Model 1; Table 3) young adult cognitive reserve [F (1,141) = 7.85, p=.006], early midlife lifestyle composite [F (1,141) = 12.93, p < 0.001], and the reserve-by-lifestyle interaction [F (1,141) = 5.40, p = 0.022] were associated with late midlife PBAD. Participants with lower young adult cognitive reserve or fewer early midlife favorable lifestyle behaviors had a predicted brain age that was older than expected given their chronological age. These results remained after adjustment for all covariates (Table 3, Model 2): cognitive reserve [F (1,104)=6.88, p=0.01)], lifestyle composite [F(1,104)=111.37, p=0.001)], reserve-by-lifestyle interaction [F(1,104)=5.24, p=0.02)]. With the exception of age, no covariates were significant in the full model.

**Table 3.** Linear Mixed Effects Models Predicting the Brain Age Difference Score (PBAD) in Late Midlife.

| Effect | Basic Model (Model 1) |  |  |  | Full Model (Model 2) |  |  |  |
| --- | --- | --- | --- | --- | --- | --- | --- | --- |
|  | Estimate<br>(SE) | t Value | Pr > \|t\| | 95%CI<br>(lower, upper) | Estimate<br>(SE) | t Value | Pr > \|t\| | 95%CI<br>(lower, upper) |
| Intercept | -33.82 (7.48) | -4.52 | <.0001 | -48.5342, -19.1022 | -46.34 (12.05) | -3.85 | 0.0002 | -70.0698, -22.6025 |
| Cognitive reserve (Age 20) | 3.40 (1.21) | 2.80 | 0.006 | 0.9995, 5.7868 | 3.57 (1.36) | 2.62 | 0.010 | 0.8739, 6.2847 |
| Modifiable lifestyle behaviors (Age 40) | 1.32 (0.37) | 3.60 | 0.001 | 0.5929, 2.0414 | 1.42 (0.42) | 3.37 | 0.001 | 0.5834, 2.2488 |
| Cognitive reserve by lifestyle interaction | -1.13 (0.49) | -2.32 | 0.022 | -2.0908, -0.1688 | -1.25 (0.55) | -2.29 | 0.024 | -2.3296, -0.1668 |
| Ethnicity(Ref=non-White) | 1.82 (0.95) | 1.91 | 0.059 | -0.06865, 3.7065 | 1.66 (1.18) | 1.40 | 0.164 | -0.6864, 4.0050 |
| Age at MRI | 0.41 (0.11) | 3.65 | 0.001 | 0.1860, 0.6255 | 0.45 (0.13) | 3.56 | 0.001 | 0.1999, 0.7019 |
| Stroke (Ref=stroke) | 1.60 (1.15) | 1.39 | 0.167 | -0.6762, 3.8802 | 0.96 (1.40) | 0.69 | 0.494 | -1.8200, 3.7468 |
| **Early Midlife Covariates:** |  |  |  |  |  |  |  |  |
| Cardiovascular index |  |  |  |  | -1.02 (1.68) | -0.61 | 0.545 | -4.3458, 2.3099 |
| Obesity (BMI>=30) |  |  |  |  | -0.64 ( 1.24) | -0.51 | 0.609 | -3.0935, 1.8227 |
| Hypertension (yes/no) |  |  |  |  | 0.95 (1.06) | 1.15 | 0.254 | -0.6585, 2.4688 |
| Diabetes (yes/no) |  |  |  |  | 0.56 (2.79) | 0.20 | 0.840 | -4.9616, 6.0908 |
| Respiratory index |  |  |  |  | -0.59 ( 1.24) | -0.47 | 0.637 | -3.0467, 1.8710 |
| Alcohol hepatitis/<br>Cirrhosis (yes/no) |  |  |  |  | 1.48 (4.66) | 0.32 | 0.752 | -7.7669,10.7175 |
| Depression (yes/no) |  |  |  |  | 0.91 (0.79) | 0.90 | 0.372 | -1.1479, 3.0402 |
| Height |  |  |  |  | 0.13 (0.09) | 1.40 | 0.166 | -0.05411, 0.311 |
| APOE (Ref=e4+) |  |  |  |  | -0.42 (0.71) | -0.60 | 0.552 | -1.8375, 0.9881 |

Abbreviations: *APOE*= Apolipoprotein E (e4+ indicates presence of any e4 allele). Model 1 N=431; df=1,141; Model 2 N=354; df=1,104 (N lower due to missing covariates). All models adjust for the random effect of family.

The reserve-by-lifestyle composite interaction for Model 2 is depicted in Figure 1. For ease of illustration, we created 2 lifestyle groups (0,1 = unfavorable; 1,2 = favorable). Higher young adult cognitive reserve (~>.75 SD above the sample mean) was protective against effects of unfavorable lifestyle on PBAD. For those with low-to-moderate young adult cognitive reserve, unfavorable lifestyles at median age 41 were associated with older predicted brain age compared to their counterparts with favorable lifestyles. The difference between these lifestyle groups tended to be greater with lower reserve.

**Figure 1.**
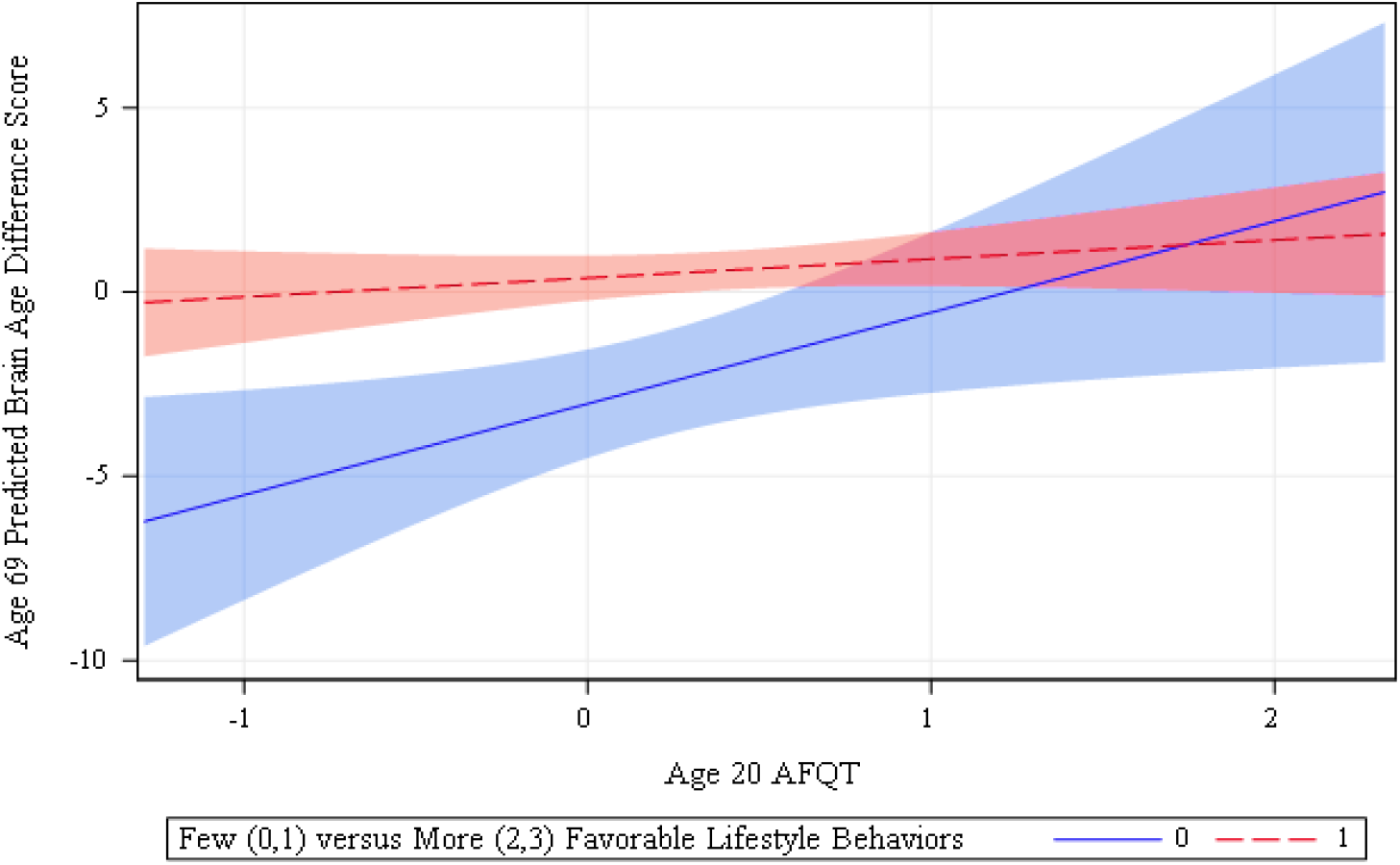
Interaction between Young Adult Cognitive Reserve and Early Midlife Modifiable Lifestyle Behaviors on Predicted Brain Age Difference Scores (PBAD) in Late Midlife. The reserve-by-lifestyle interaction is illustrated by categorizing lifestyle into two groups low (0,1) versus more (2,3) favorable lifestyle behaviors assessed at median age 41. Cognitive reserve is the continuous measure of cognitive ability assessed at median age 20.

Having an older than predicted brain age was associated with MCI at median age 69 (F (2,130) = 3.13; p = 0.05). The non-amnestic MCI group had older than predicted brain age (mean PBAD = −3.62, SE = 1.14) compared with the amnestic MCI group (mean PBAD = −0.61, SE = 1.06; t = −2.31; p = .02; 95%CI −5.578,-0.431) or cognitively normal group (mean PBAD=-1.15, SE =.73; t=-2.37; p=.02; 95%CI −4.535, −0.406).

## Discussion

Young adult cognitive reserve and modifiable lifestyle behaviors in early midlife predicted brain aging over a period spanning four decades. High cognitive reserve appeared to be protective regardless of lifestyle behaviors, but favorable lifestyle was protective for those with low-to-moderate cognitive reserve. Consistent with studies of much older adults, predicted brain age was associated with MCI.^19,22^ In those studies brain age also predicted conversion to AD. AD patients’ brain age appears older than their chronological age by about 10 years.^18^ In the much younger VETSA sample, participants with non-amnestic MCI had older-than-predicted brains than either the amnestic MCI or cognitively normal groups. Favorable lifestyle behaviors such as not smoking or low alcohol consumption are presumed to contribute to global brain integrity and cognitive health through multiple pathways involving cardiovascular risk and neurotoxic effects^48–52^. Mechanisms behind social integration’s benefits are less clear.^6,53,54^ These results suggest that unfavorable lifestyle behaviors as early as midlife increase risk for dementia, but it remains unclear of if the effect is related to AD *per se*.

Our study is unique in its examination of the cognitive reserve-by-lifestyle interaction. There is evidence that overall cognitive ability peaks in the 20s-to-early 30s, so these participants may have been close to peak reserve at the time of the AFQT assessment.^35^ In addition, the lifestyle composite data were collected earlier in the life course compared with most studies of long-term effects of lifestyle on the brain.^1,2^ These results may be thought of as indicating that, regardless of level of young adult cognitive reserve, favorable lifestyle confers resistance to advanced brain aging regardless; however, high reserve confers both resistance to advanced brain aging and resilience against the effects of toxic lifestyle behaviors.^13,55^ PBAD also suggests brain maintenance,^56^ but confirmation of brain maintenance requires longitudinal brain data.

Meta-analyses suggest that a brain health risk reduction agenda could be effective in reducing risk for dementia.^2,3,57^ Both the U.K. National Institute of Health and Care Excellence, and the U.S. National Institutes of Health list smoking and social integration as among the top modifiable risk factors for dementia. The 2020 Lancet Commission reported that a third of dementia incidence can be attributed to modifiable risk factors^1^, although this remains to be fully supported by clinical trials.^58^ Smoking, alcohol consumption, and social integration in the first half of the life cycle may be potential modifiable targets for early intervention for brain health. In one randomized controlled trial, for example, brain volume increased across 40 weeks in an intervention involving social interaction.^54^

There are several limitations to this study. Participants were all men and mostly non-Hispanic White, thus results may not generalize across genders or different racial or ethnic origins. Lifestyle indicators were self-reported; however, that is typical for such measures. We also cannot definitively rule out whether PBAD was already different at the time lifestyle behaviors were assessed since we did not have a measure of predicted brain age at that time. Because patterns of atrophy for global brain aging might be different from the patterns associated with AD,^59^ further follow-up will be needed to determine the extent to which PBAD may be a longitudinal risk factor for AD or MCI.

### Conclusions

Given the large number of persons likely to develop dementia in the next decades, early identification of modifiable risk factors may be key to identifying and reducing the incidence and burden of AD and other dementias. These results highlight the potential value of focusing on preventive efforts earlier in development and not just in later life. Given the evidence suggesting that favorable lifestyle confers resistance to advanced brain aging particularly among those with lower cognitive reserve, efforts at prevention could be prioritized among those with lower cognitive reserve. With increasing number of individuals with MCI and AD combined with a lack of effective treatments, early risk reduction and prevention can be major components in efforts to facilitate maintenance of brain integrity, thereby mitigating individual and societal health, economic, and social burden.

## Acknowledgements

The content is the responsibility of the authors and does not necessarily represent official views of the NIA, NIH, or VA. The U.S. Department of Veterans Affairs, Department of Defense; National Personnel Records Center, National Archives and Records Administration; National Opinion Research Center; National Research Council, National Academy of Sciences; and the Institute for Survey Research, Temple University provided invaluable assistance in the creation of the VET Registry. The Cooperative Studies Program of the U.S. Department of Veterans Affairs provided financial support for development and maintenance of the Vietnam Era Twin Registry. We would also like to acknowledge the continued cooperation and participation of the members of the VET Registry and their families. Ms. Warren was supported by the Advancing Diversity in Aging Research (ADAR) program at San Diego State University (R25 AG043364).

## Online supplemental tables

**eTable 1.**
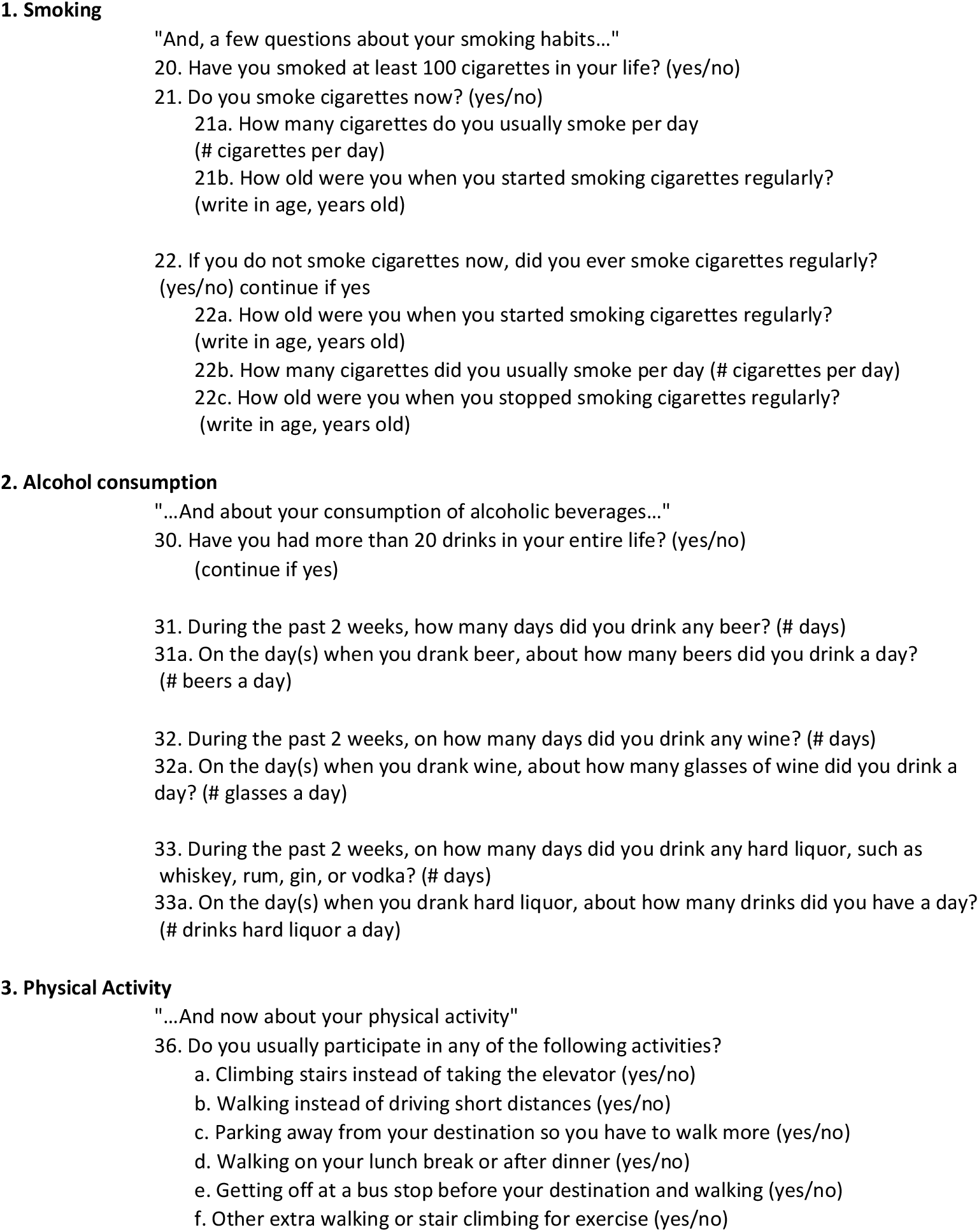
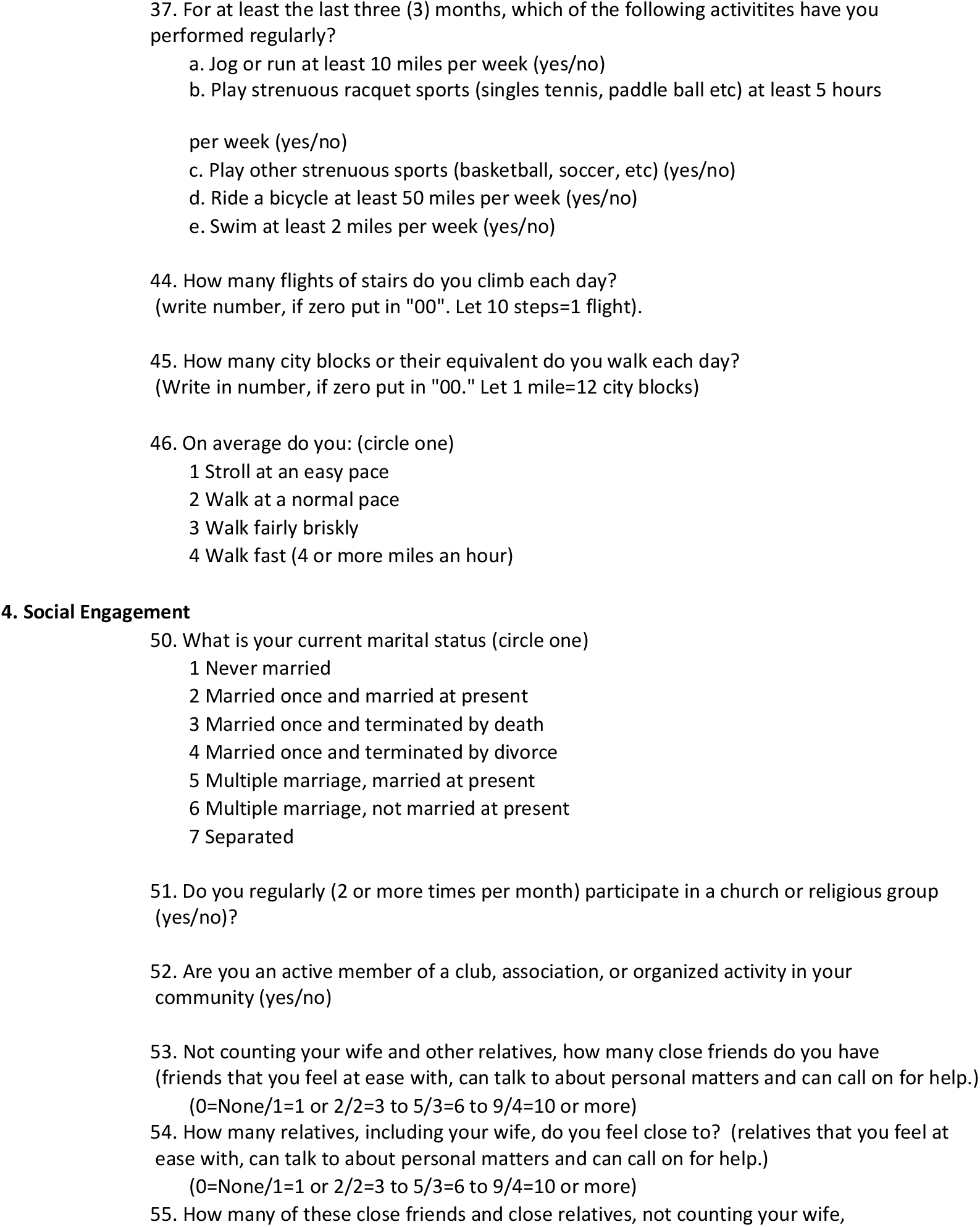
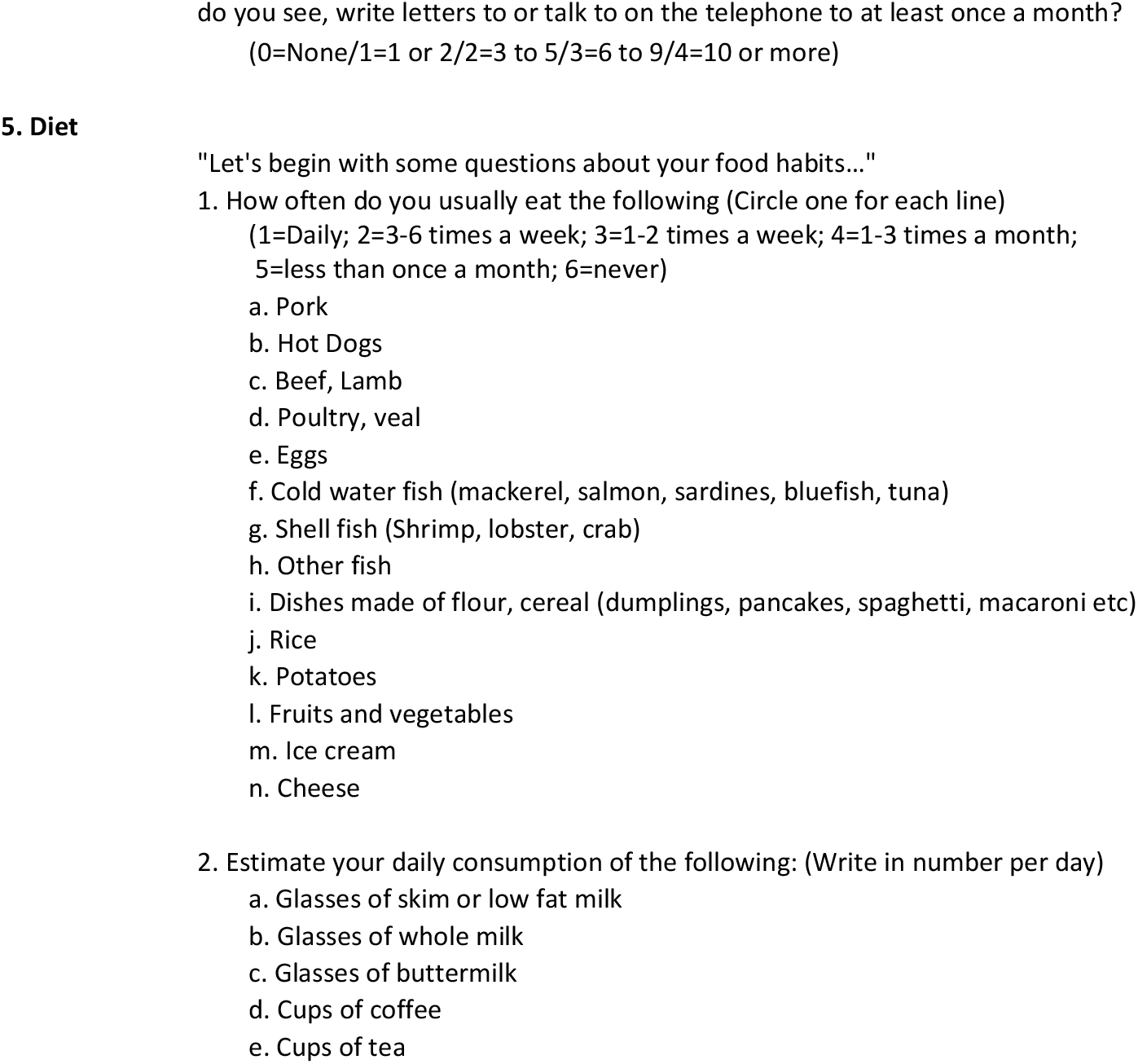
Items for Modifiable Lifestyle Behaviors from Median Age 41 Data Collection.

**eTable 2.** Frequencies of Early Midlife (Median Age 41) Covariates.

|  |  | Frequency<br>(0=no/1=yes) | Percent |
| --- | --- | --- | --- |
| Cardiovascular index | 0 | 443 | 97.79 |
|  | 1 | 10 | 2.21 |
| Depression | 0 | 416 | 92.44 |
|  | 1 | 34 | 7.56 |
| Diabetes | 0 | 455 | 99.34 |
|  | 1 | 3 | 0.66 |
| Hypertension | 0 | 385 | 84.62 |
|  | 1 | 70 | 15.38 |
| Liver Disease Index | 0 | 449 | 99.78 |
|  | 1 | 1 | 0.22 |
| Obesity (BMI >= 30) | 0 | 433 | 94.75 |
|  | 1 | 24 | 5.25 |
| Respiratory Index | 0 | 423 | 94.84 |
|  | 1 | 23 | 5.16 |

Note: 1=presence of disease/condition 0=absence of disease/condition; BMI=body mass index

**eTable 3.** Correlations between Predictor Variables and Covariates.

|  | PBAD | Early Adult<br>Cognitive<br>Reserve | Modifiable<br>Lifestyle<br>Composite |
| --- | --- | --- | --- |
| Age 20 AFQT (cognitive reserve) | 0.192 |  |  |
| p-value | <.0001 |  |  |
| Modifiable Lifestyle Behavior Composite | 0.241 | 0.171 |  |
| p-value | <.0001 | 0.000 |  |
| Education (lifetime max) | 0.155 | 0.345 | 0.266 |
| p-value | 0.001 | <.0001 | <.0001 |
| Ethnicity (1=non-Hispanic White/0=Other) | -0.181 | -0.339 | -0.086 |
| p-value | <.0001 | <.0001 | 0.073 |
| Stroke (VETSA 3; 1=yes/0=no) | -0.138 | -0.074 | -0.168 |
| p-value | 0.002 | 0.100 | 0.000 |
| **Early Midlife (Age 40) covariates** |  |  |  |
| Liver Disease (1=yes/0=no) | -0.025 | 0.037 | -0.024 |
| p-value | 0.599 | 0.439 | 0.615 |
| Respiratory Problems (1=yes/0=no) | 0.019 | 0.052 | -0.086 |
| p-value | 0.686 | 0.279 | 0.076 |
| Cardiovascular Problems (1=yes/0=no) | -0.041 | -0.043 | -0.030 |
| p-value | 0.381 | 0.361 | 0.532 |
| Depression (1=yes/0=no) | -0.012 | -0.077 | -0.121 |
| p-value | 0.806 | 0.107 | 0.012 |
| Diabetes (1=yes/0=no) | -0.027 | -0.022 | -0.042 |
| p-value | 0.571 | 0.644 | 0.380 |
| Obesity (1=BMI >=30/0=BMI < 30) | 0.028 | 0.049 | -0.021 |
| p-value | 0.551 | 0.304 | 0.666 |
| High Blood Pressure (1=yes/0=no) | -0.065 | -0.051 | -0.010 |
| p-value | 0.165 | 0.280 | 0.829 |
| Height (inches) | 0.101 | 0.088 | 0.020 |
| p-value | 0.023 | 0.050 | 0.675 |

Note: PBAD=Predicted Brain Age Difference Score; BMI=Body Mass Index

**Supplemental Table 4.** Correlations among Key Variables and Lifestyle Behaviors.

|  | PBAD | Lifestyle Composite | Early Adult Cognitive Reserve |
| --- | --- | --- | --- |
| Stroke (by late midlife) | -0.138 | -0.168 | -0.074 |
| (p-value) | 0.002 | 0.000 | 0.100 |
| Early Midlife Lifestyle Behaviors |  |  |  |
| Diet | -0.017 | 0.064 | 0.125 |
| (p-value) | 0.719 | 0.190 | 0.009 |
| Smoking | 0.138 | 0.695 | 0.232 |
| (p-value) | 0.003 | <.0001 | <.0001 |
| Alcohol Consumption | 0.192 | 0.525 | -0.023 |
| (p-value) | <.0001 | <.0001 | 0.636 |
| Physical Activity | -0.073 | 0.034 | 0.035 |
| (p-value) | 0.126 | 0.479 | 0.465 |
| Social Engagement | 0.127 | 0.582 | 0.069 |
| (p-value) | 0.007 | <.0001 | 0.146 |

Note: PBAD=Predicted Brain Age Difference Score. For lifestyle behaviors, higher scores indicate more favorable behaviors.

